# A histidine switch regulates pH-dependent filament formation by the caspase-9 CARD

**DOI:** 10.1101/2025.06.02.657148

**Authors:** Swasti Rawal, Ðesika Kolarić, Stefan Bohn, Dagmar Kolb, Tea Pavkov-Keller, Iva Pritišanac, Tobias Madl, Ambroise Desfosses, T. Reid Alderson

**Affiliations:** Research Unit Integrative Structural Biology, Medicinal Chemistry, Otto Loewi Research Center, Medical University of Graz, 8010 Graz, Austria; Helmholtz Munich, Molecular Targets and Therapeutics Center, Institute of Structural Biology, 85764 Neuherberg, Germany; Technical University of Munich, TUM School of Natural Sciences, Department of Bioscience, Bavarian NMR Center, 85747 Garching, Germany; Helmholtz Munich Cryo-Electron Microscopy Platform, 85764 Neuherberg, Germany; Core Facility Ultrastructure Analysis, Center for Medical Research, Gottfried Schatz Research Center, Medical University of Graz, 8010 Graz, Austria; Institute of Molecular Biosciences, University of Graz, 8010 Graz, Austria; BioTechMed-Graz, 8010 Graz, Austria; Institut de Biologie Structurale, Université Grenoble Alpes, CEA, CNRS, IBS, 38000 Grenoble, France

**Keywords:** death domain folds, caspase activation and recruitment domain (CARD), helical filament, cryo-EM, NMR

## Abstract

The caspase activation and recruitment domain (CARD) mediates protein-protein interactions in apoptotic and inflammatory signaling pathways. In humans, more than 30 proteins contain a CARD, several of which have been reported to polymerize into helical filaments. Here we found that the CARD from the apoptotic protease caspase-9 (C9^CARD^) self assembles into filaments *in vitro* at physiological pH and salt concentrations. The C9^CARD^ more readily polymerizes under low-salt or mildly acidic conditions, suggesting a significant role for electrostatic interactions in mediating filament formation. Using NMR spectroscopy, we determined the p*K*_a_ of the lone histidine residue, H38, which supports a role for histidine protonation in enhancing filament formation. Indeed, mutation of H38 to introduce a positive (H38R) or negative (H38D) charge, or to remove the pH-dependence of the side chain at this site altogether (H38N), dramatically alters the filament propensity of the domain. Using cryo-election microscopy, we determined 3.4- and 3.2-Å structures of the wild-type and H38R C9^CARD^ filaments, respectively, which provide new insights into the molecular basis of C9^CARD^ polymerization and its pH dependence via H38.

## Introduction

Apoptosis, or programmed cell death, is an intricate biochemical signaling cascade that eliminates damaged cells and aids in the regulation of cellular differentiation and homeostasis^1–3^. Dysregulated apoptosis plays a central role in unregulated inflammatory responses, neurodegenerative diseases, and cancer development^4–7^. Caspase-9 (C9), a cysteine-aspartic protease^8,9^, plays a key role as an initiator of the intrinsic apoptotic pathway^10,11^ and regulates cell death processes by cleaving and activating downstream effector caspases, such as C3 and C7 ^12–14^. Under basal conditions, C9 is present as an inactive monomer and requires clustering and activation by the apoptosome^10,15,16^, a heptameric scaffold comprising Apaf-1, ATP, and cytochrome C. While the role of C9 in activating effector caspases is well established, it is increasingly clear that C9 cleaves hundreds of substrates with diverse implications^17,18^.

At the molecular level, C9 is a 416-residue protein that is comprised of a caspase activation and recruitment domain (CARD, residues 1-96), an interdomain linker, and a protease domain (residues 139-416). Activation of the protease domain involves substrate-driven, proximity-induced dimerization on the apoptosome platform^19,20^ . Protein engineering approaches have further demonstrated that C9 can be activated via recruitment to synthetic platforms or by constitutive dimerization of the protease domain^21^. The dimerization-dependent activity of C9 was leveraged as a safety switch in adoptive T-cell therapy, whereby the expression of a chimeric protein comprising the protease domain of C9 and a chemically-inducible dimerization domain enables targeted and rational control over C9 activation and cell death^22^. Moreover, the activation or catalytic activity of C9 can be inhibited allosterically by post-translational modifications that are distal from the active site ^23–28^, as well as by zinc and peptide binding to other allosteric sites on the protease domain^29,30^. More generally, allostery in caspase protease domains appears to be widespread ^31–38^, including in C9 where it displays half-of-sites reactivity ^19,20^, which is an extreme form of negative cooperativity.

The N-terminal CARD domain belongs to the death-domain fold (DDF) superfamily, which fold into conserved six-helix bundles and mediate protein-protein interactions^39–41^. The DDF superfamily comprises four sub-families that include the CARD, death domain (DD), death effector domain (DED), and pyrin domain (PYD). DDFs primarily make homotypic interactions within sub-families (CARD-CARD, PYD-PYD, etc.) with remarkable specificity^40^. In C9, its CARD (C9^CARD^) mediates recruitment to the apoptosome platform via CARD-CARD interactions with Apaf-1^CARD^. The CARD-CARD complex forms a helical oligomer or disk ^15,42–46^, and the same helical-disk architecture involving C9^CARD^ and Apaf-1^CARD^ has been observed in cryo-electron microscopy (EM) structures of the apoptosome:caspase-9 complex^47,48^. In full-length C9, the CARD appears to tumble independently of the protease domain^20^, although transient inter-domain interactions may influence substrate recruitment^49^. CARDs often use multiple interfaces to interact with themselves and others, including in C9 where the C9^CARD^:Apaf-1^CARD^ complex involves three different interfaces^43,45,48^. The locally asymmetric binding interfaces among CARD complexes can lead to complex modes of homo- and hetero-oligomerization, such as polymerization into helical filaments^50–58^ and the formation of biomolecular condensates via liquid-liquid phase separation^59^. For caspases, the formation of supramolecular complexes via DDF self-assembly dramatically increases the local concentration of their protease domains, leading to rapid activation. For example, filament formation was shown to play a role in the activation of the CARD-containing C1^51^ as well as the tandem DED-containing C8^60,61^.

Here, we report that the C9^CARD^ polymerizes into filaments *in vitro* and present a 3.2-Å cryo-EM structure of the C9^CARD^ filament. We dissected the biophysical determinants of C9^CARD^ self-assembly, and by manipulating the pH or salt concentration we identified conditions that enhance C9^CARD^ filament formation. We identified a histidine “switch” (H38) whose protonation stimulates filament assembly, and we found that the mutation H38R enhances filament propensity. We subsequently determined a 3.4-Å cryo-EM structure of the H38R variant of C9^CARD^, which serves as a mimic of the histidine-protonated form. Other mutations at this position to introduce a negative charge (H38D) or non-titratable polar group (H38N) decrease filament propensity, suggesting that H38 may function as a pH sensor that regulates filament assembly.

## Results

We expressed and purified recombinant, ^15^N-labeled labeled C9^CARD^ for NMR spectroscopy (**Figure 1A**). The two-dimensional ^1^H-^15^N heteronuclear single-quantum coherence (HSQC) spectrum of the ^15^N-labeled C9^CARD^ at pH 6.5 and room temperature confirmed that our construct is folded under these conditions with the expected number of backbone amide signals. We also observed an additional 10 signals from aliased Arg Nε-Hε groups, which contain ^15^N chemical shifts near 85 ppm, as confirmed by increasing the ^15^N spectral width (**Supplementary Figure 1**). Next, we recorded triple-resonance NMR data on a ^13^C,^15^N- labeled sample and assigned the backbone, Cβ, and Arg Nε-Hε resonances. The secondary ^13^Cα chemical shifts indicate that the six helices are formed in solution (**Figure 1B**), including a kink in helix-1. This is further confirmed by TALOS-N-derived secondary structure determination (**Supplementary Figure 2**). ^1^H^N^ temperature coefficients suggest that the secondary structure elements are stably hydrogen-bonded in solution (**Figure 1C**), although possible fraying of helices may be occurring at the N- or C-terminal ends of helix-1 and helix-2. Moreover, ^15^N spin relaxation experiments confirm that the C9^CARD^ is monomeric in solution under these conditions with a rotational correlation time of approximately 6.5 ns (**Figure 1C**). This falls between an empirical value of 7 ns for a protein of this molecular mass^62^ (from a standard curve obtained 5 °C lower than our conditions), as well as the value predicted by HydroNMR (5.5 ns) using a crystal structure of C9^CARD^ that lacks six residues found in our construct. To test for transient oligomerization that may slightly increase the rotational correlation time, we also measured ^15^N Carr-Purcell-Meiboom-Gill relaxation dispersion experiments in the range of 20-35 °C; however, we did not detect any millisecond-exchange contributions to the ^15^N transverse relaxation rate (**Supplementary Figure 2**). Thus, the C9^CARD^ is folded and monomeric under these conditions, giving rise to high-quality NMR data in solution.

**Figure 1.**
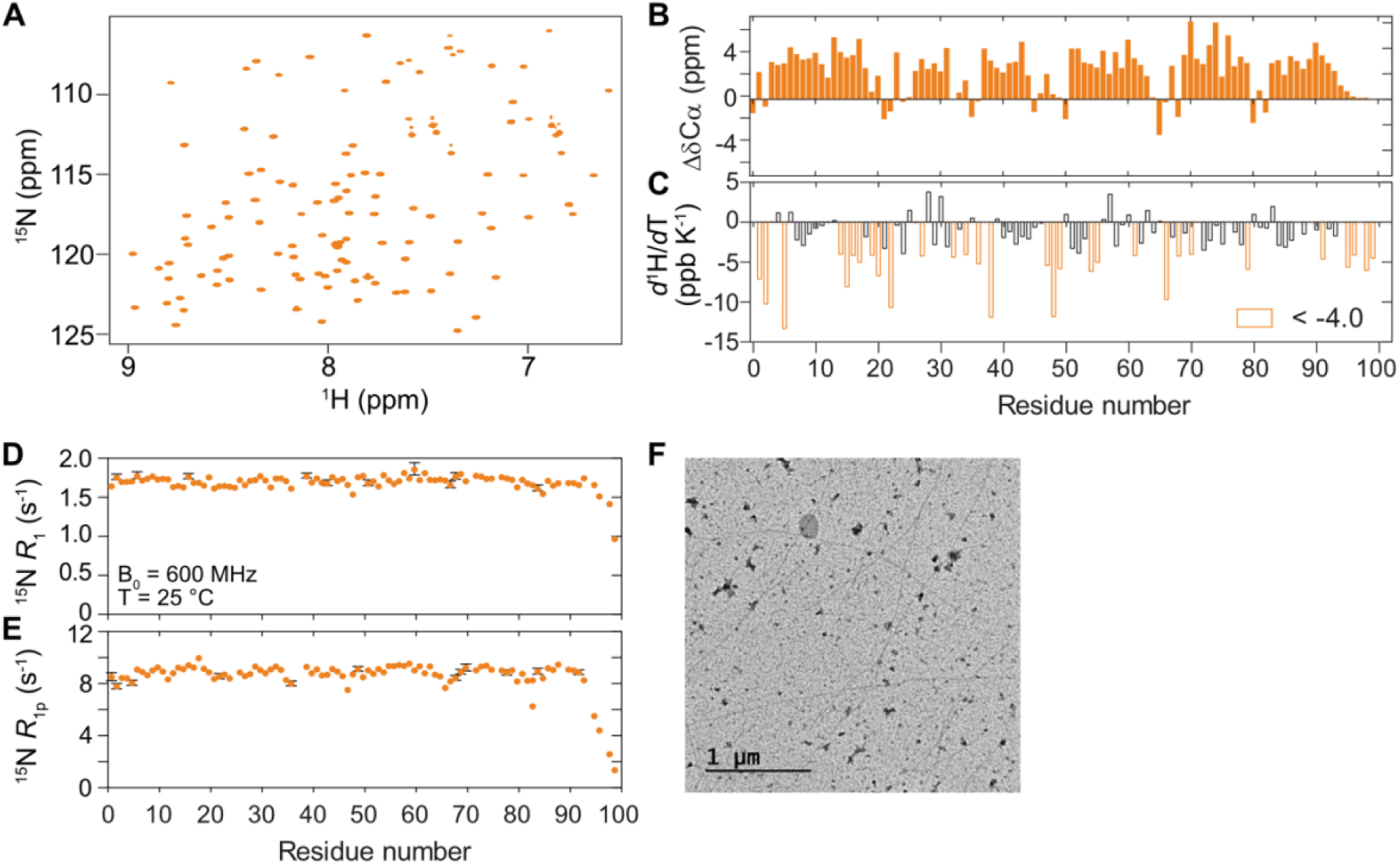
C9^CARD^ polymerizes from a monomeric state into filaments. (**A**) 2D ^1^H-^15^N HSQC spectrum of uniformly ^15^N-labeled C9^CARD^ in 25 mM HEPES, 50 mM NaCl, 5 mM TCEP, 0.5 mM EDTA at pH 6.5 and 25 °C. (**B**) Secondary ^13^Cα chemical shifts (Δδ^13^Cα) and (**C**) ^1^H^N^ temperature coefficients (*d*^1^H/*d*T) for the C9^CARD^. Residues with *d*^1^H/*d*T values below -4.0 ppb/K, suggestive of non-hydrogen-bonded amides, are shown in orange. (**D**) ^15^N *R*_1_ and (**E**) ^15^N *R*_1ρ_ relaxation rates recorded at 25 °C and a static magnetic field strength of 14.1 T (600 MHz). The ^15^N spin-lock field strength used for the *R*_1ρ_ experiment was 2 kHz. (**F**) Representative negative-stain EM image of C9^CARD^ filaments that were obtained from a sample in the absence of salt at pH 5.5 where the maximum solubility of C9^CARD^ is 150 μM (*vide infra*).

### C9^CARD^ self-assembles into filaments

During the NMR sample preparation, we noticed that C9^CARD^ aggregated above protein concentrations of *ca.* 0.3 mM in salt-free buffers. Interestingly, the addition of 500 mM NaCl to the C9^CARD^ increased its solubility to above 3 mM protein concentration (**Figure 1D**), suggesting that salt ions screen out electrostatic interactions between C9^CARD^ molecules that otherwise cause aggregation. Indeed, despite having an expected net charge of zero at pH 6.5, the primary structure of C9^CARD^ is significantly enriched in charged residues, with nearly 33% of the sequence comprising R, K, D, and E, and equal numbers of positively charged (15 Arg, 1 Lys) and negatively charged residues (9 Asp, 7 Glu) (**Supplementary Figure 3**).

We next sought to test the hypothesis that the aggregates of C9^CARD^ are filaments. To this end, we used negative-stain EM to examine the properties of aggregates that were obtained from a sample of C9^CARD^ in salt-free buffer at pH 7. We observed filament-like structures with an average diameter of approximately 10 nm in the negative-stain EM micrographs (**Figure 1E**).

### Protonation of H38 promotes filament assembly

Having established that the C9^CARD^ forms filaments, as observed by nsEM, we next sought to find solution conditions that promote its filament-forming propensity. To this end, we screened the effects of pH and salt concentration on the stability and solubility of the C9^CARD^. Notably, the thermodynamic stability of the C9^CARD^ was markedly reduced as the pH was lowered from 7 to 5.5 (**Figure 2A**), with a concomitant decrease in solubility (**Figure 2B**). We confirmed by nsEM that the aggregates that formed under acidic conditions were also filamentous (**Supplementary Figure 4**). Since these experiments used only the isolated CARD domain, we tested whether inclusion of the interdomain linker (residues 100-138) and the protease domain (residues 139-416) impacted filament formation. We purified the C9^CARD+linker^ construct (residues 1-138) and full-length C9 (1-416) harboring an inactivating C287A mutation that prevents auto-proteolysis of the p20-p10 linker ^63^. As compared to the isolated CARD domain, both the C9^CARD+linker^ and full-length C9^C287A^ were less prone to aggregate (**Figure 2B**). We collected negative-stain EM data to visualize the aggregates formed by these longer constructs, finding that the majority of aggregates were amorphous, although several filaments could be identified (**Supplementary Figure 5**). Thus, we proceeded to structurally characterize the isolated C9^CARD^ filaments henceforth.

**Figure 2.**
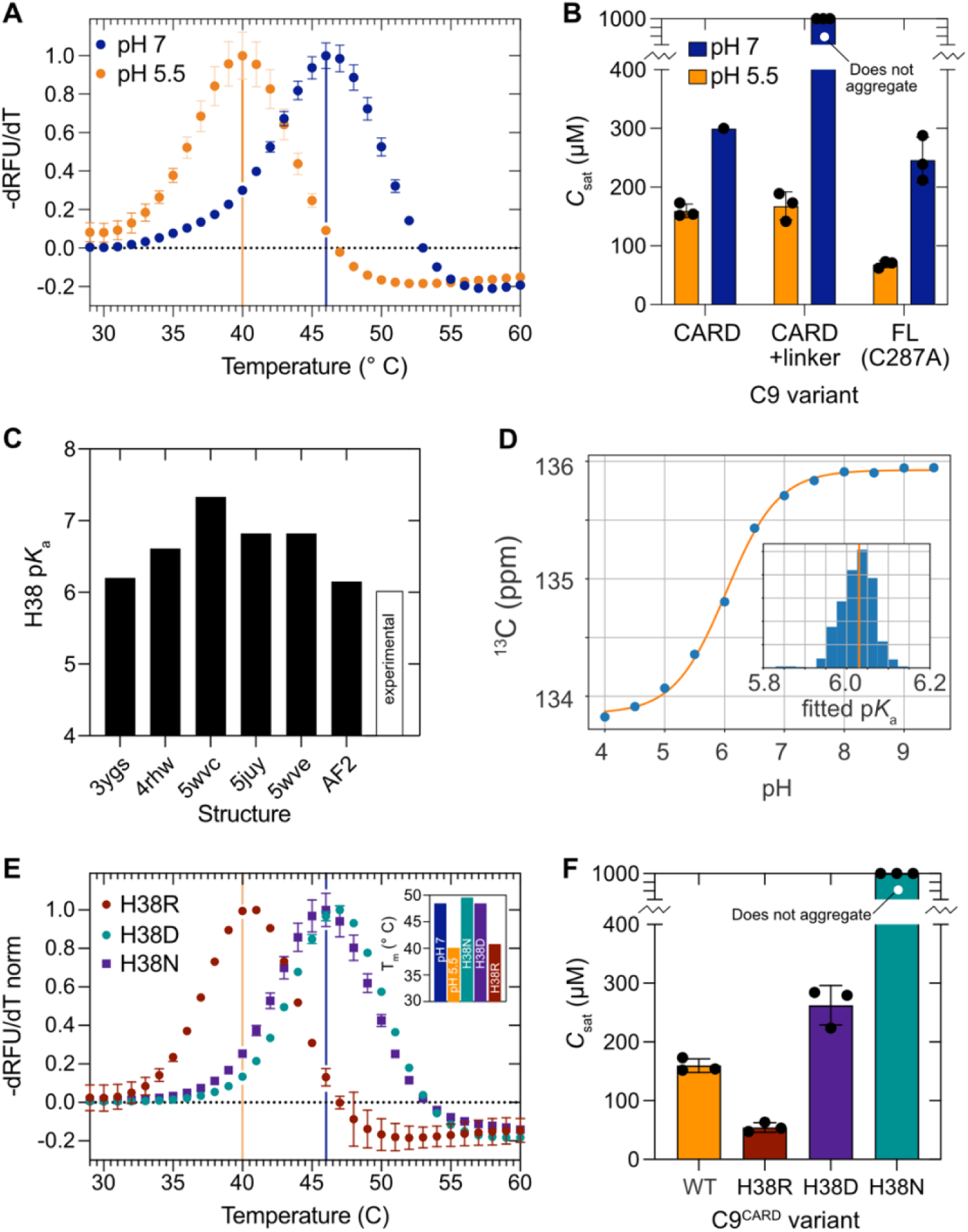
H38 protonation and the H38R mutation destabilize the CARD and enhance filament assembly. (**A**) DSF-based thermal melt of C9^CARD^ in 25 mM HEPES, 2 mM DTT at pH 7 (blue) or pH 5.5 (orange). The normalized, negative derivative of the change in relative fluorescence units (RFU) with temperature is shown on the y-axis. (**B**) *C*_sat_ measurements for different variants of C9 at pH 7 (blue) or pH 5.5 (orange). The buffer is the same as panel A. (**C**) Structure-based prediction of H38’s p*K*_a_ as a function of the input structure (black, filled). AF2 corresponds to C9 model deposited the AlphaFold Protein Structure Database. The NMR-derived, experimentally determined p*K*_a_ value for H38 is shown in white. (**D**) Variation in the H38 ^13^Cε chemical shift as a function of the pH, with the solid line corresponding to a fit to the Henderson-Hasselbalch equation. The inset shows the distribution of p*K*_a_ values obtained from a bootstrap error analysis, with the orange line positioned at the mean value of 6.03. (**E**) DSF-based thermal melt of C9^CARD^ variants H38R (red), H38D (green), and H38N (purple). The curves are compared to the WT C9^CARD^ at pH 7 (blue line) or pH 5.5 (orange line). The inset depicts the derived melting temperatures for each variant. (**F**) *C*_sat_ determination for the H38 variants at pH 5.5 and 25 °C. The H38R variant is less soluble than the wild-type CARD.

Our observations suggest that the protonation of one or more titratable side chains destabilizes the C9^CARD^ at pH 5.5 and enhances its propensity to self-assemble into filaments. Among its titratable side chains, the C9^CARD^ contains a single histidine residue (H38) with a theoretical p*K*_a_ of 6.5 that falls within the pH range described above (**Figure 2C**). However, p*K*_a_ values can be dramatically shifted away from their theoretical values by local interactions, including electrostatics^64–66^. For this reason, we first used PROPKA^67^ to perform C9^CARD^ structure-based p*K*_a_ prediction (PDB/chain: 4rhw/E, 3ygs/P, 5wvc/B, AlphaFoldDB/A, 5juy/O, 5wve/S), which returned a mean H38 p*K*_a_ value of 6.7 ± 0.4 (6.61, 6.20, 7.33, 6.15, 6.82, and 6.82, respectively) (**Figure 2C**). We then experimentally determined the p*K*_a_ of H38 via NMR spectroscopy to monitor pH-induced changes to the ^13^Cε chemical shift of H38 (**Figure 2D**). Quantitative analysis revealed an apparent p*K*_a_ of 6.03 ± 0.04 (**Figure 2D**), which indicates that H38 is, indeed, predominantly protonated at pH 5.5 where C9^CARD^ exhibits enhanced filament propensity. Given the large number of acidic side chains in C9^CARD^ (9 Asp, 7 Glu), it is also likely that one or more sites are titrating in the pH range below 5.

### H38R mutation mimics the low-pH state and triggers filament assembly

Given that H38 is mostly protonated at pH 5.5 where the C9^CARD^ is highly filamentous, we next evaluated the impact of altering the electrostatic properties at this position while pH remains neutral. To this end, we created three variants of the C9^CARD^ in which we altered the residue at position 38 to be positively charged (H38R), negatively charged (H38D), or polar but not titratable (H38N) (**Figure 2E**). We then measured the melting temperature and solubility of each variant at neutral pH. We observed that the H38R variant (T_m_ = 41 °C) was destabilized relative to the wild-type C9^CARD^ (47 °C) as well as the H38D (47 °C) and H38N (49°C) variants (**Figure 2E**). Indeed, the melting temperature of the H38R variant at pH 7 resembles that of the wild-type C9^CARD^ at pH 5.5, which further indicates that a positive charge at this position is destabilizing at neutral pH. By contrast, the H38D and H38N variants melt at the same or higher temperature as the wild-type C9^CARD^ at neutral pH.

We observed similar trends with protein solubility: under identical solution conditions (25 mM MES, 2 mM DTT, pH 5.5), the H38R variant aggregated at a nearly 6-fold lower protein concentration as compared to the wild-type CARD (**Figure 2F, Supplementary Figure 7**). Under these conditions, H38 in the wild-type C9^CARD^ is predominantly, but not fully protonated (*ca.* 25%/75% un/protonated at pH 5.5 with p*K*_a_ = 6.03). The six-fold lower *C*_sat_ of H38R suggests that the positive charge of the Arg head group, with an expected p*K*_a_ above 10, plays a role in polymerization. By contrast, H38D and H38N remained soluble up to protein concentrations that were respectively two- and more than 20-fold higher than the wild-type C9^CARD^ (**Figure 2F**), with no aggregates of H38N detected in this experiment.

Using nsEM, we observed filaments of H38R that resembled those formed by the wild-type C9^CARD^ with an average diameter near 10 nm (**Supplementary Figure 7**). Similarly, we observed filaments for the H38D variant that only formed at significantly higher protein concentrations as compared to the H38R variant (**Supplementary Figure 8**), suggesting that altering the residue at this site modulates but does not inhibit filament formation. Collectively, our results indicate that a positive charge at position 38 thermodynamically destabilizes the CARD, lowers its solubility, and promotes self-assembly into the filamentous state. These results support the notion that H38 is the primary pH sensor in the range of 7 to 5.5, where its protonation (or H38R mutation) contributes to polymerization.

Next, we analyzed the variation in residue type at this position of the CARD in more than 250 different C9 orthologs. The most common residue type at this position is the negatively charged Asp (*n* = 179), followed by A (16), P (14), T (12), L (10), H (7), N (7), G (6), and S (6) (**Supplementary Figure 8**). We did not find a positively charged residue at this position in any of the examined organisms. In agreement with our biophysical results above, these bioinformatics data indicate that positively charged residues are not preferred at this position in C9^CARD^.

### H38 protonation alters its side-chain dynamics and clashes with the helix dipole

H38 is a solvent-exposed residue in the C9^CARD^ that is located close to two negatively charged residues (E41, D42). It is, therefore, unclear why the H38R mutation and protonation of H38 significantly destabilize the domain. Indeed, the structure-based predictors of protein stability FoldX^68^ and DDmut^69,70^ do not report any significant changes to the stability of the C9^CARD^ following the H38R mutation (ΔΔG = -0.19 kcal mol^-1^ vs. RMSE of 1.37 kcal/mol on cross-validation set). This contrasts with our experimental observations (**Figure 2**).

Thus, to better understand why H38 protonation and H38R are destabilizing, we performed all-atom molecular dynamics (MD) simulations of the C9^CARD^ as a function of the H38 protonation state or with the H38R mutation. In the crystal structure of C9^CARD^ (PDB: 4rhw, chain E), which was crystallized at pH 5.5, the side chain of H38 is surrounded by nearby charged residues in helix-3, forming interactions with the side chains of E41 (3.6 Å) and D42 (3.7 Å). The acidic pH of the crystallization buffer suggests that H38 was likely protonated. For the C9^CARD^ with H38 in either the neutral or protonated form, our MD simulations show that the global fold of the domain is preserved with a mean backbone and side chain root-mean-square deviation (RMSD) values near 2 and 3 Å, respectively, over the course of the trajectory (**Supplementary Figure 9**). For most backbone atoms, per-residue root-mean-square fluctuation (RMSF) values indicate negligible differences (± 0.1 Å) for the majority of sites in C9^CARD^ (**Supplementary Figure 9**). In the protonated form, locally elevated backbone RMSF values are only observed at the N-terminus for residues M1 (H38 neutral – H38 protonated = -1.0 Å) and D2 (-0.3 Å), at a few sites in helix-1 (residues 11-13 and 16; -0.2 to -0.3 Å), in loop 4,5 (65-68, -0.2 to -0.8 Å), and at the C-terminus (94,95; -0.3 Å). On the other hand, locally decreased RMSF values in protonated-H38 are seen in loop 3,4 (residues 44-46, 47; 0.15-to-0.2 Å) near H38 itself (**Supplementary Figure 9**). For H38 itself, there is no difference in backbone RMSF for the neutral or protonated state. However, among side-chain heavy-atom RMSF values, H38 shows the second largest difference in RMSF values (**Supplementary Figure 9**) (neutral – protonated = - 0.5 Å), pointing to increased structural dynamics in the protonated form. In addition, the sites discussed above with non-zero differences in backbone RMSF values similarly exhibit non-zero differences in side-chain RMSF values (**Supplementary Figure 9**).

Consistent with the above RMSD and RMSF analyses, we found that H38 protonation does not cause any significant changes to the backbone phi and psi dihedral angle distributions that were observed in the H38-neutral state. Instead, we observed large redistributions to some side-chain rotamer populations, emanating from H38 itself (**Figure 3A, 3B, 3C**). In its neutral form, H38 almost exclusively populates the *gauche+* rotamer (*ca.* 98% population), whereas its protonation triggers complete conversion to the *trans* (48%) and *gauche-* (52%) rotamers without impacting its backbone dihedrals (**Figure 3D**).

**Figure 3.**
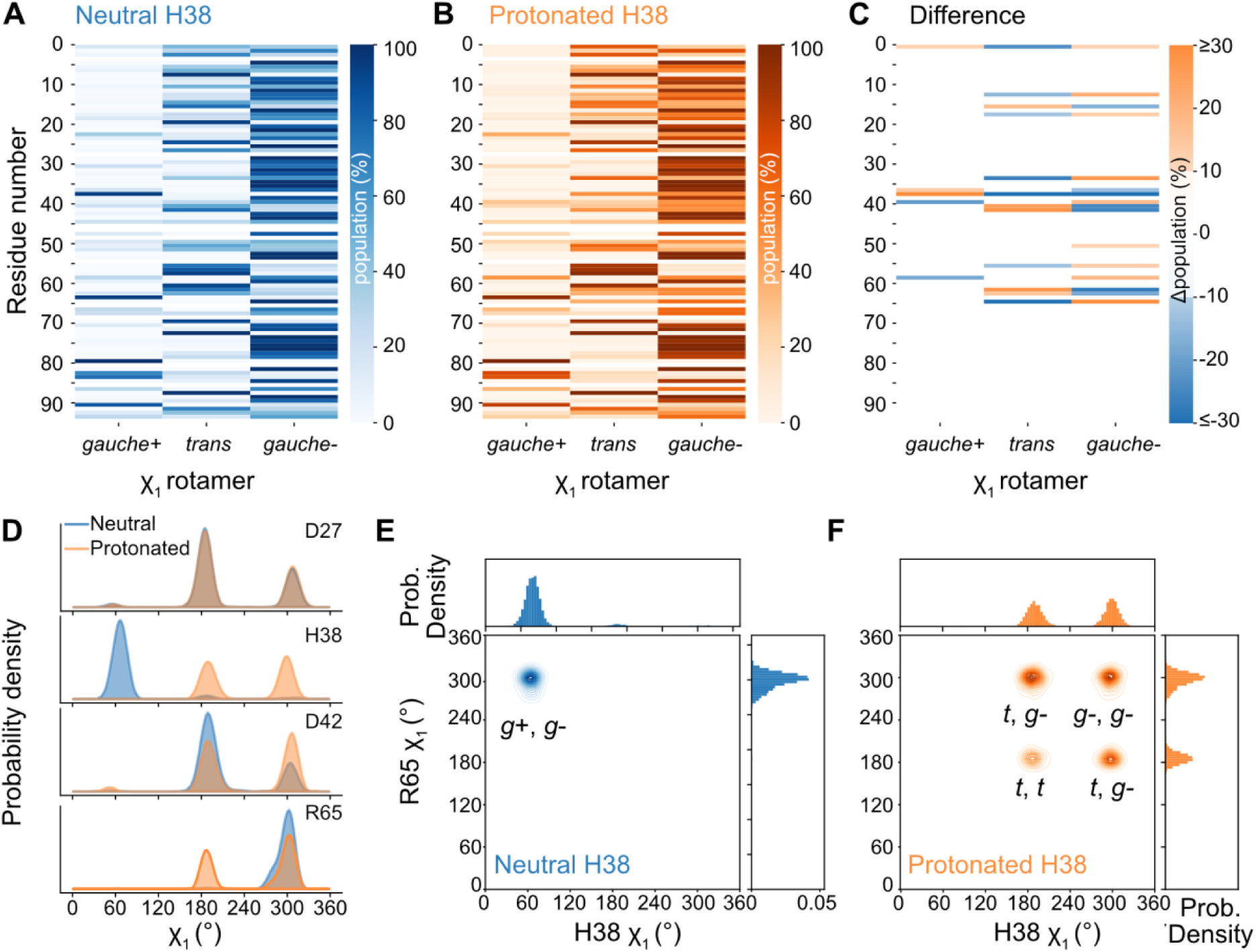
Protonation of H38 leads to a redistribution of side-chain χ_1_ torsion angles. χ_1_ rotamer populations (*gauche*+, *trans*, *gauche-*) derived from the 5-μs MD simulations of the H38-neutral (**A**) or H38-protonated (**B**) states of the C9^CARD^. Residues with empty data points correspond to Ala and Gly residues. (**C**) Population difference plot (protonated minus neutral), where orange (blue) values are positive (negative) and correspond to an enriched (depleted) rotamer in the H38-protonated state. (**D**) Rotamer distributions over the trajectories for H38-neutral (blue) and H38-protonated (orange) for the indicated side chains. (**E**, **F**) Two-dimensional surfaces showing the increased conformational heterogeneity upon H38 protonation for the side chains of H38 and R65.

The origin of this rotameric change likely involves the helix dipole at the N-terminus of helix-3, where H38 is located as an N-cap residue. In its neutral form, the imidazole ring of H38 points inward toward the dipole of helix-3, perhaps stabilizing the partial positive charge of the helix dipole. Upon protonation of the imidazole ring, however, the positive charges would experience an electrostatic clash; thus, the *trans* and *gauche-* rotameric states of H38 position the imidazole ring away and nearly perpendicular to the helix dipole, thereby minimizing a direct clash. Furthermore, our measured p*K*_a_ value of 6.03 for H38 agrees with studies in model peptides where a His residue in an N-cap position yields a p*K*_a_ of 6.1 as compared to 7.4 in a C-cap position^70–72^ . In addition, the protonation of a His residue in an N-cap position significantly destabilizes helical peptides^73^, which agrees with our thermal melting data above.

### Cryo-EM structures of the C9^CARD^ filament and its H38R variant

To examine the structural properties of the C9^CARD^ filament in more detail, we determined high-resolution structures of the WT and H38R CARD filaments with cryo-EM (**Figure 4**). For the WT filament, we observed bundle formation in which multiple filaments clustered together. We were able to overcome the bundles in our cryo-EM data analysis, eventually obtaining a 3.2-Å map. By contrast, the H38R filaments were long and straight, without significant bundle formation, leading to determination of a slightly lower resolution 3.4- Å model. Our cryo-EM derived models of the C9^CARD^ subunit are highly similar to the available crystal structures of the C9^CARD^, with backbone root-mean-square deviation values near 0.4 Å (**Figure 4**), showing that the overall tertiary structure is preserved within the filament. Both filaments display *C*_2_ symmetry with left-handed, two-start helical assembly consisting of approximately five subunits per turn. The H38R filament has a twist angle of -67.2° and an axial rise of 9.1 Å per subunit. The inner and outer diameters of the WT and H38R filaments are respectively 1.7 and 9.2 and 1.5 and 9 nm (**Figure 4A, 4B**).

**Figure 4.**
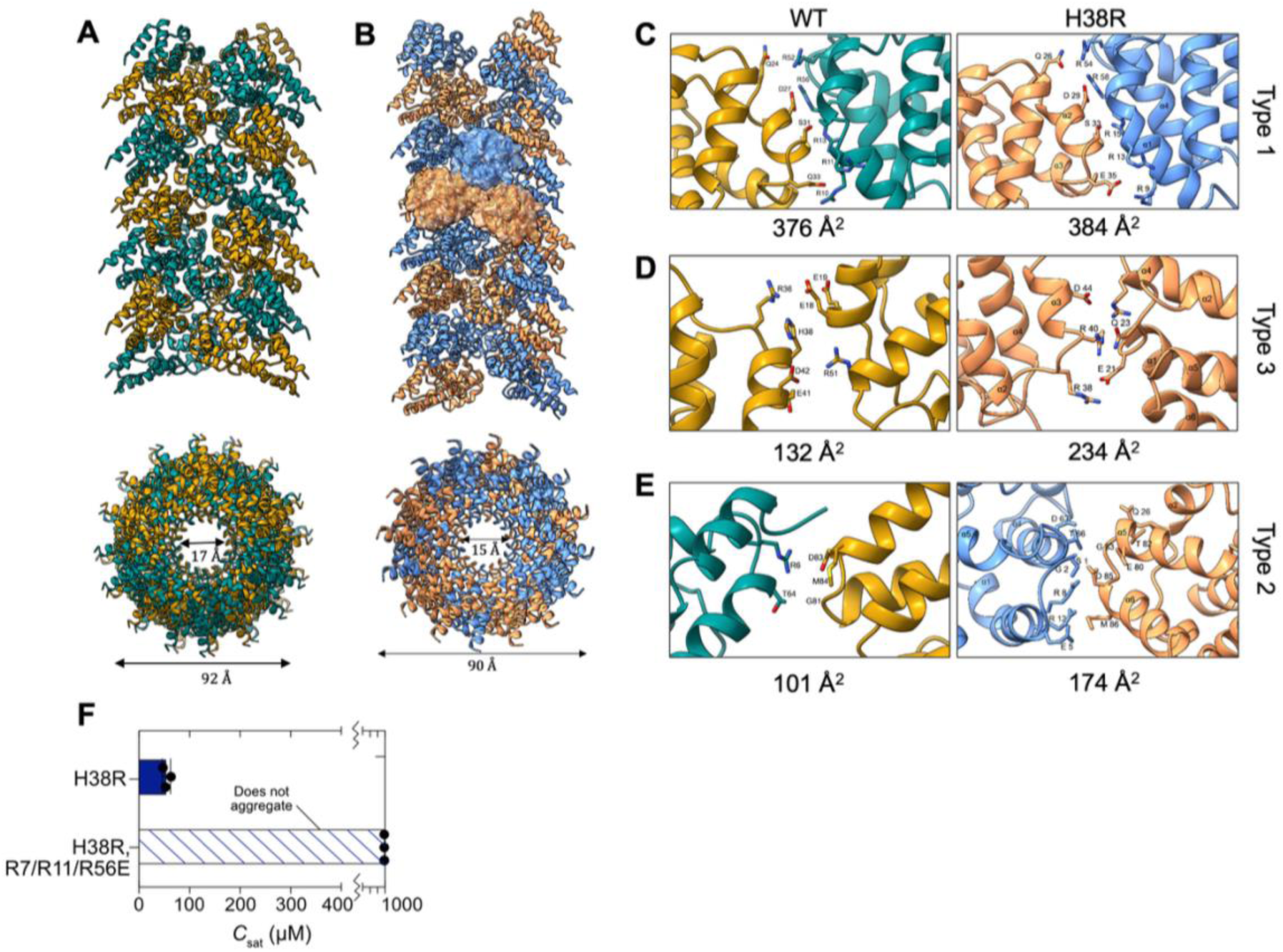
Cryo-EM structures of the C9^CARD^ filament and its H38R variant. (**A**) Side view of the cryo-EM structure of the C9^CARD^ filament, including a rotated view down the filament that depicts the inner and outer diameters. (**B**) The same as in A but for the H38R filament. Zoomed-in regions that correspond to the type-1 (**C**), type-3 (**D**), and type-2 (**E**) interfaces are shown along with the corresponding interface area in square Å. (**F**) A type-1 interface triple mutant introduced into the H38R variant (R7E/R11E/R56E) is prevents filament formation. The *C*_sat_ could not be determined, because the triple mutant did not aggregate in this assay.

Similar to other DDF filaments, the C9^CARD^ filament involves three major asymmetric interfaces with opposing interacting surfaces designated as *a* and *b* (**Figure 4C, 4D, 4E**). The WT and H38R filaments share similar interfaces, with the type-1 and -2 interfaces mediating intra-strand interactions between the helical turns, whereas the type-3 interface forms inter-strand interactions in the helical strand direction. In the H38R filament, the type-2 and type-3 interfaces bury slightly more area than the in the WT filament. H38 is located in the type-2 interface, and thus its mutation to Arg increases the interaction surface area. The type-1 interface is the most extensive due to electrostatic complementarity, covering a total buried surface area of 376 Å^2^ and 384 Å^2^ in the WT and H38R filaments (**Figure 4C**), and thus likely contributes the most to the binding energy required for filament formation. This interface mediates interaction between residues in the α2 and α3 helices (e.g., E33, S31, D27, Q24) of one subunit with residues in the α1 and α4 helices (e.g., R52, R56, R13, R11, R7) of the neighboring subunit. Within a distance of 4 Å, there are a total of six hydrogen bonds and four salt bridges contributing to this interface. The type-3 interface (**Figure 4D**) buries 132 Å^2^ and 234 Å^2^ of surface area, respectively, contains H38 as well as salt bridges between the guanidinium group of arginine and side chains of glutamate and aspartate. In total, there are four salt bridges and three hydrogen bonds that contribute to this interface. The type-3 interface mediates interaction between residues in the α2-α3 loop and the α3 helix of one subunit with the residues in the α4 helix and α1-α2 loop of the neighboring subunit. The type-2 interface, with a total buried surface area of 101 Å^2^ and 174 Å^2^, respectively (**Figure 4E**), is the least extensive among all three interfaces. It constitutes only two hydrogen bonds between residues in the α5-α6 loop and α6 helix of one chain to the α1 helix of the neighboring chain.

To test the cryo-EM model of the C9^CARD^ filament, we generated a triple-mutant in which the mutations R7E, R11E, R56E were introduced to the H38R variant (**Figure 4F**). These charge-reversing mutations were designed to disrupt interactions across the type-1 interface of the C9^CARD^ filament. We purified the H38R/triple-mutant and measured its solubility under filament-forming conditions at pH 5.5. However, we were not able to induce aggregates, suggesting that it is unable to form filaments even at high protein concentrations above 3 mM. Thus, the structure-based mutations that we introduced into the H38R variant raised its *C*_sat_ from *ca.* 50 μM to over 3 mM, corresponding to an increase of more than 60-fold. Finally, we compared the molecular interactions within the C9^CARD^ filament to those formed with the Apaf-1 CARD (Apaf^CARD^) in the apoptosome. The type-1b interface of the filament involves residues in the α1 and α4 helices, which form the primary interaction surface for the Apaf^CARD^ (PDB: 3ygs), suggesting that the C9^CARD^ filament would be unable to associate with Apaf^CARD^ or the apoptosome.

### Comparing CARDs from the paralogs C9 and C1

Given the observation that the C9^CARD^ self-assembles into filaments, and with access to its high-resolution structure, we next compared the sequence and structure of C9^CARD^ to its paralog C1^74,75^. C1 mediates homotypic CARD-CARD interactions and self-assembles into filaments during the innate immune response^76,77^. Structural alignment of the helices in C1^CARD^ (PDB: 5fna) and C9^CARD^ (PDB: 4hrw) yields a backbone root-mean-square deviation of 1.6 Å, indicating highly similar structures. We analyzed the per-residue sequence conservation among C1 and C9 orthologs (**Figure 5A**), from which it is evident that buried residues have a higher degree of sequence conservation, as expected given the structural restraints imposed by the three-dimensional fold of the CARD. Indeed, when we quantified the relative solvent accessible surface area (rSASA), we observed that buried residues are more likely to be conserved in both C9 and C1 orthologs (**Figure 5B, 5C**). This result is consistent with previous work that quantified the rate of sequence changes in surface residues versus buried residues^78,79^.

**Figure 5.**
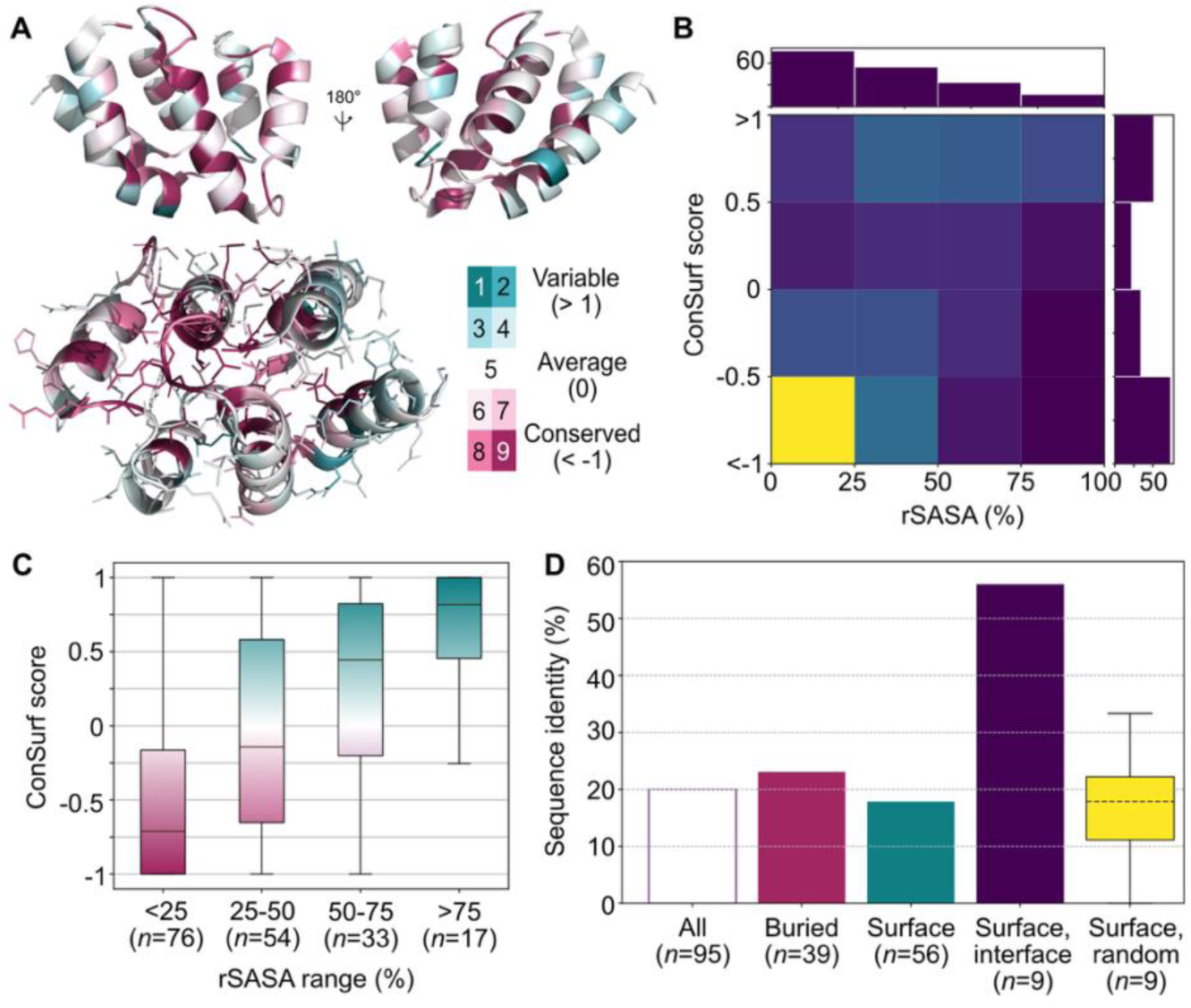
Comparison of C9^CARD^ and C1^CARD^ sequences. (**A**) ConSurfDB^81^-derived sequence identity among C9 orthologs mapped onto the structure of the C9^CARD^ (PDB: 4rhw). The degree of conservation is shown with the color bar. Amino-acid side chains show that buried residues appear more conserved than those on the surface. (**B**) 2D histogram showing the per-residue relative solvent accessible surface area (rSASA) versus the ConSurf^81^ score for both the C9^CARD^ and C1^CARD^. (**C**) Box-plot representation of the data in panel B, plotted here as a function of the rSASA range in 25%-increment steps between 0-100%. The number of residues in each bin is listed below. The boxes include data points between the first and third quartiles, with the whiskers extending to 1.5-fold the interquartile range. (**D**) Pairwise sequence identity between C1^CARD^ and C9^CARD^ computed for all residues in the alignment (All), only buried residues (pink, rSASA < 25%), only surface residues (teal, rSASA ≥ 25%), or only surface residues involved in the type-1 filament interface (purple). The number of residues in each is shown below. A box plot of the distribution of pairwise sequence identities between randomly sampled surface residues (*n* = 9) over 10,000 simulations is shown in yellow.

A global pairwise sequence alignment of the two CARDs reveals only 20% sequence identity (**Figure 5D**). When considering only buried (surface) residues, the pairwise sequence identity only slightly increases (decreases) (**Figure 5D**), indicating that the sequence differences are widespread. However, detailed inspection of the C1^CARD^ and C9^CARD^ surfaces reveals additional insight into their evolutionary relationships. When considering only the surface residues that comprise the largest type-1 interface in the C1^CARD^ filament^51^ (PDB: 5fna, R10, K11, R15, N23, D27, E28, Q31, D52, R55), we find that the C9^CARD^ and C1^CARD^ exhibit 56% identity at these nine positions (**Figure 5D**). To test the significance of this value, we randomly sampled surface residues (*n* = 9) from the C1^CARD^ 10,000 times and quantified the pairwise sequence identity to C9^CARD^ at these sites. This random-sampling approach yielded a mean sequence identity of 18 ± 12% among surface residues, which corresponds to a Z-score of 3.2 and an empirical p value of 1 × 10^-4^, indicating that the type-1 interface residues on the surface are statistically more conserved than random chance. This observation is surprising because surface residues typically evolve at a four-fold faster rate than buried residues in the hydrophobic core of proteins^78,79^. On the other hand, conserved surface residues can signify shared and functionally relevant protein-protein interaction sites; yet the C1^CARD^ and C9^CARD^ do not share any known DDF interactors^80^. An alternative explanation is that the conserved surface residues are involved in a homo-oligomeric interface, like the type-1 filament interface of these filaments.

## Discussion

CARDs are protein-protein interaction modules that function in apoptotic and inflammatory signaling pathways^39,41,82^. Even though CARD domains have high structural similarity, their activation mechanisms and assembly architectures vary and range from monomers, dimers, soluble oligomers, or insoluble filaments^41^. The C9^CARD^ has served as a model system for structural and biophysical studies of CARDs, with multiple high-resolution structures available as well as insight into its oligomeric architecture that forms upon binding to Apaf-1^CARD^ in the apoptosome ^10,15,16^. Here, we found that the C9^CARD^ can access yet another structural form and polymerizes into filaments *in vitro* at physiological pH and salt concentrations. The propensity to form filaments was enhanced under low-salt or acidic conditions, suggesting a significant role for electrostatic interactions in driving filament formation. Charge-altering mutations of the lone histidine residue to introduce a positive (H38R) or negative (H38D) charge, or to remove the charge at this site altogether (H38N), dramatically modulated the self-assembly of the C9^CARD^, indicating that interactions involving H38 contribute to filament formation.

Using NMR spectroscopy, we determined the p*K*_a_ of H38 to be 6.03 ± 0.04, confirming that the side chain is predominantly protonated under acidic conditions at pH 5.5 where the C9^CARD^ is highly filamentous. We further employed an integrative biophysical and structural approach to understand the destabilizing effect of H38 protonation or the H38R mutation, identifying an electrostatic clash with the partial positive charge from the helix dipole. We used cryo-electron microscopy to determine 3.2- and 3.4-Å structures of the WT C9^CARD^ and H38R filaments. Like other DDFs filament structures, C9^CARD^ shows extensive electrostatic interactions between the CARD subunits within the filament at the type-1 interface, while H38 is involved in the type-3 interface and the H38R increases the size of this interface.

The paralogous C1, which shares the same domain architecture with C9, is known to form filaments during its activation^51^. Paralogs arise through gene duplication and often retain structural similarity; however, functional specialization driven by evolutionary pressures can lead to substantial sequence divergence^83–85^. Indeed, at the sequence level, C1^CARD^ and C9^CARD^ have highly diverged, with only 20% identity between the two domains. Nonetheless, we found that the surface residues that comprise the type-1 filament interface are over 50% identical between the two paralogs, which far exceeds expectations from random sampling of the solvent-accessible surface. This observation suggests that residues at the type-1 interface may have been more preserved after functional divergence.

In context of full-length C9, filament formation by the CARD would lead to an increase in the local effective concentration of the attached protease domain that may prime it for catalytic activity. Activation of the protease domain requires dimerization, which very weakly dimerizes (*K*_d_ > 10 mM) in the absence of substrate^20^. For four C9 molecules that were tethered to the apoptosome, the local effective concentration (*C*_eff_) was determined to be 0.5 mM^20^, which is far below the dimerization *K*_d_ and insufficient to trigger activation in the absence of substrate. On the other hand, in the filamentous state, the *C*_eff_ of the protease domain is expected to be significantly higher due to the larger number of molecules involved in the filament, which could lead to auto-activation of C9 in an analogous way to C1^51^.

Our experiments in this study were performed *in vitro* with purified components, and thus it remains unclear if the C9^CARD^ self-assembles into filaments *in vivo*. However, the relatively high *C*_sat_ concentration of the C9^CARD^ at physiological pH (*ca.* 150 μM), which was obtained in the absence of salt, suggests that a “priming” event may be required to nucleate C9^CARD^ filaments *in vivo*. Given that the concentration of C9 is expected to be in the nanomolar range in human cells^86^, filament formation would require clustering of C9 molecules to reach the significantly higher *C*_eff_. As mentioned above, in the case of C9 binding to the apoptosome, the *C*_eff_ of the protease domain approaches 500 μM, despite its tethering via a long and disordered linker ^20^. The *C*_eff_ of the C9^CARD^ on the apoptosome, by definition, would be significantly larger due to the smaller volume occupied by the four C9 molecules; however, the C9 residues involved in the filamentous interface would be occupied by interactions with Apaf-1^CARD^. Thus, filaments involving C9^CARD^ would only be possible upon dissociation from Apaf-1^CARD^ and clustering, or upon a nucleation or priming event that presents available and exposed C9^CARD^ molecules for further polymerization.

Interestingly, Borgeaud and colleagues^87^ recently reported that, in cells, the apoptosome forms large, transient assemblies or foci that depend on the Apaf-1^CARD^. The apoptosome foci did not form in cells that were devoid of C9, suggesting a potential link between foci formation and CARD-CARD interactions involving Apaf-1 and C9 ^87^. Similarly, the CARD from C2 was recently reported to be necessary for liquid-liquid phase separation in the presence of ubiquitin^59^, further highlighting the range of assemblies that are accessible to caspase CARDs in cells. While the physiological relevance of the C9^CARD^ filament remains to be tested, we

In summary, we identified a helical filament that is formed by the C9^CARD^ and investigated the biophysical properties that drive filament formation. We identified a histidine residue (H38) whose protonation or Arg-mutation stimulates filament assembly, thus functioning as a pH-sensitive switch that regulates CARD polymerization. Indeed, removing the charge at this position via the H38N mutation generated a highly soluble protein whose *C*_sat_ could not be determined. Cryo-EM structures of the wild-type and H38R C9^CARD^ filaments reveal a similar overall architecture to other CARD filaments and expand the overall catalog of known DDF filaments. Our identification of a natural, pH-responsive protein filament may be relevant for the *de-novo* design of protein filaments: Shen and colleagues have recently described an approach to design pH-responsive protein filaments that leverage His protonation^88^. Taken together, our findings suggest that C9^CARD^ polymerization is finely tuned by local physicochemical cues, particularly pH, through a single histidine residue that modulates charge at a structurally sensitive site.

## Methods

### Protein expression and purification

DNA encoding for human caspase-9 (UniProt ID: P55211) was synthesized and codon optimized for *E. coli* expression by GenScript^20^. The gene was then sub-cloned into a pET-29b(+) expression vector that contained an N-terminal hexahistidine-tag followed by small ubiquitin-like modifier (His-SUMO) solubility tag. Two fragments of caspase-9 were derived from the full-length plasmid via mutagenesis: the CARD domain (residues 1-99) and the CARD+linker (residues 1-138)^20^. All mutagenesis, including point mutants of H38 (vide infra) and elsewhere in the CARD, was performed by GenScript. All of the caspase-9 expression constructs contain a Ser-Gly linker that separates His-SUMO from Met1 of the CARD domain. This was found to be necessary to enable complete cleavage of the His-SUMO tag by the Ulp1R3 SUMO protease^20^, likely for steric reasons due to the burial of Met1 in the structure of the CARD.

For recombinant protein expression and purification, we followed previously established protocols for C9^CARD^+linker and C9 (C287A) ^20^. All variants were expressed and purified similarly. For the production of ^15^N-labeled or ^13^C,^15^N-labeled protein, M9 minimal medium was used with 1 g/L of ^15^NH4Cl and 6 g/L of D-glucose or 1 g/L of NH_4_Cl and 2 g/L of uniformly ^13^C-labeled D-glucose, respectively. Protein purification proceeded as outlined above.

### NMR Spectroscopy

All NMR samples were prepared in in 5-mm NMR tubes with 5% ^2^H_2_O added for the lock. To obtain resonance assignments of the C9^CARD^, a ^13^C,^15^N-labelled sample of C9^CARD^ was prepared at a protein concentration of 1.4 mM in 25 mM MES at pH 6.5 with 50 mM NaCl. Backbone (^1^H^N^, ^1^H^α^, ^15^N, ^13^CO, ^13^Cα) and side-chain (aliphatic ^13^C/^1^H, Arg ^15^Nε/^1^Hε) resonances were assigned based on three-dimensional (3D) HNCO, HN(CA)CO, HNCA, HN(CO)CA, HNCACB, C(CO)NH, H(CCO)NH spectra^89^ recorded at 20 °C on a Bruker Avance-III 700-MHz spectrometer equipped with a cryogenically cooled, 5-mm TCI probe. Non-uniform sampling was employed in which the data sampling density in the indirect dimensions ranged from 10-25% and was based on a random sampling schedule. Spectral reconstruction was performed using SMILE^90^. All spectra were processed using NMRPipe^91^ and analyzed using NMRFAM-Sparky^92^. For secondary structure determination, the assigned backbone and ^13^Cβ chemical shifts were analyzed using TALOS-N^93^.

^15^N longitudinal (*T*_1_) and transverse rotating-frame (*T*_1ρ_) spin relaxation experiments^94^ were collected on ^15^N-labeled C9^CARD^ at a protein concentration of 0.84 mM in 25 mM HEPES, 50 mM NaCl, 2 mM DTT at pH 6.5 with the temperature set to 25 °C. Data were recorded on a 600-MHz Bruker Avance Neo NMR spectrometer with a room-temperature probe. The pulse sequences employed HSQC-based detection with sensitivity enhancement^95^. Magnetization was sampled at delay times of 0.01, 0.1, 0.25, 0.38, 0.5 (x2), 0.62, and 0.75 seconds, with a duplicated value at 0.5 seconds for error analysis. In the *T*_1ρ_ experiment, a 2-kHz ^15^N spin-lock field was applied to suppress chemical exchange during the transverse relaxation delay time (T), with ^1^H inversion pulses applied at T/4 and 3T/4 intervals to suppress cross-correlated relaxation between ^1^H-^15^N dipolar coupling and ^15^N chemical shift anisotropy ^96^ Magnetization was sampled at delay times of 5, 15, 30 (x2), 50, 75, 100, and 150 ms, with the 30-ms time point duplicated for error analysis. In both experiments, 8 scans were collected per FID with an inter-scan delay of 3 seconds. The ^15^N and ^1^H spectral widths were 1095 and 9615 Hz with 50* and 615* complex points collected in each dimension, respectively corresponding to acquisition times of 45.6 and 64 ms. The ^15^N spin relaxation datasets were processed with NMRPipe and NMRFAM-Sparky was used to pick peaks for subsequent lineshape fitting in FuDA^97^.

For the H38 p*K*_a_ determination, a ^13^C,^15^N-labeled C9^CARD^ sample was prepared, and the pH was varied in 0.5-unit steps between 4 and 9.5 via the addition of concentrated HCl or NaOH aliquots. For each pH value, a 2D ^1^H-^13^C HSQC spectrum was recorded, and we focused on the His ^13^Cε chemical shift given that this site is sensitive to protonation of the imidazole ring but not impacted by the relative populations of the two different neutral tautomers (δ, ε) ^98–101^ . The variation in the ^13^Cε chemical shift of H38 was fit to the Henderson-Hasselbalch equation:

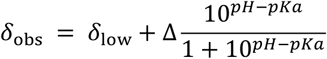

Where *ẟ*_obs_ is the observed chemical shift, *δ*_low_ the measured chemical shifts at pH 4, and Δ is the difference in chemical shift between the neutral and protonated forms. Least-squares fitting was performed in Python using the *lmfit* package ^102^, and the error was estimated from a standard bootstrap with 1,000 iterations.

### HydroNMR simulation

The C9^CARD^ crystal structure (4rhw, chain E) was supplied as input to the HydroNMR^103^ software program, which was used via the NMRbox resource^104^. In the simulation, the temperature was set to 25 °C to match the ^15^N relaxation experiments, with the solvent viscosity adjusted to 1 cPa. All other parameters, including the minimum and maximum radii of beads in the shell, were left at default values. The simulations assumed an N-H bond-length of 1.02 Å and a ^15^N chemical shift anisotropy of -160 ppm. The simulated rotational diffusion tensor of the C9^CARD^ yields values for *D*_xx_, *D*_yy_, and *D*_zz_ of 2.86, 2.89, and 3.39 × 10^7^ s^-1^, respectively, yielding an isotropic rotational correlation time of *D*_iso_ = 1/[6Tr(*D*)] = 5.47 × 10^-9^ s, where Tr(*D*) corresponds to the trace of the 3×3 rotational diffusion matrix, or 1/3 (*D*_xx_ + *D*_yy_ + *D*_zz_). However, the rotational diffusion of C9^CARD^ appears approximately axially symmetric, with the anisotropic rotational diffusion values *D*_x_ = 2.90 × 10^7^ s^-1^, *D*_y_ = 2.79 × 10^7^ s^-1^, and *D*_z_ 3.43 × 10^7^ s^-1^ corresponding to values of 2*D*_z_ / (*D*_x_ + *D*_y_) = 1.20 and *D*_x_/*D*_y_ = 1.04.

### Differential Scanning Fluorimetry

To measure the melting temperature of C9^CARD^ variants, we mixed protein sample and SYPRO Orange dye (Thermo Fisher Scientific) in a 96-well plate (Bio-Rad) to a total volume of 25 µL per well. All conditions contained a protein concentration of 30 µM in 25 mM MES and 2 mM DTT at pH 5.5 or 7. SYPRO orange dye was added to each well in 1:2500 dilution, and the plate was then sealed. Fluorescence was measured in a Bio-Rad CFX96 RT-PCR instrument with the temperature increased from 20 °C to 95 °C in 0.5-°C increments every 30 seconds. The melting temperature was calculated from the first derivative of fluorescence versus temperature. All measurements were performed in triplicates, and the reported values are the mean and one standard deviation, normalized to the maximum and minimum values of each run.

### Negative-stain Electron Microscopy

A carbon-coated copper grid (400-mesh) (Science Services, Germany) was glow-discharged using a PELCO easiGlow™ Glow Discharge Cleaning System (Ted Pella Inc., USA)) to make it hydrophilic for homogenous sample distribution. 10 μL of purified C9^CARD^ (25 mM HEPES, 200 mM NaCl, 0.5 mM EDTA, 2 mM TCEP, pH 7.5) was adsorbed onto the glow-discharged grid and allowed to settle for 1 minute. The remaining sample was carefully removed with filter paper, and immediately 10 μL of a 1% (w/v) uranyl acetate staining solution was added. The uranyl acetate solution was allowed to settle for 1 minute and the excess solution was removed using filter paper. The sample was air dried for few minutes before putting inside the microscope. Sample visualization was done on a Tecnai G2 FEI (Thermo Fisher Scientific Inc., USA) microscope equipped with an Ultrascan 1000 CCD camera (Gatan AMETEK, Germany) operating at an acceleration voltage of 120 kV.

### Structure-based pK_a_ prediction

PDB files were uploaded to the PDB2PQR webserver that assigns titration states with version 3.6.1 of PROPKA^67^. The pH of the calculation was set to 7.0 and the default forcefield was used. Steric clashes were removed with the debump algorithm inside PDB2PQR that optimizes inter-atomic distances that fall below defined threshold values of 1.0 Å for hydrogen-hydrogen, 1.5 Å for hydrogen-heavy atom, and 2.0 Å for heavy atom-heavy atom collisions. Hydrogen bond networks were optimized by PDB2PQR by enabling the flipping of His, Asn, and Gln side chains, the rotation of side-chain hydrogen atoms on Ser, Thr, Tyr, and Cys residues, and placement of side-chain hydrogen atoms on His, Asp, and Glu.

### Molecular Dynamics Simulations

We performed all-atom molecular dynamics simulations with the GROMACS^105^ simulation package using the AMBER99SB-ILDN force field^106^. We selected the highest-resolution crystal structure of the C9^CARD^ (PDB: 4rhw) and extracted chain E from the PDB file. The system was solvated in a rhombic dodecahedron of TIP4P-EW^107^ water molecules with a minimal distance of 10 Å between the protein and the box edge, and then energy minimized using the steepest descent algorithm to remove any steric clashes, with the maximum force set to 1000 kJ mol^-1^ nm^-1^. Following minimization, the system was equilibrated in two steps: *NVT* equilibration at 300 K for 100 ps, using the v-rescale thermostat^108^, and *NPT* equilibration at 300 K and 1 bar for an additional 100 ps, with Berendsen pressure-coupling^109^. During equilibration, all protein heavy atoms were restrained to their initial positions. Production MD simulations were carried out in the *NPT* ensemble at 300 K and 1 bar using periodic boundary conditions. The particle mesh Ewald method^110^ was employed for long-range electrostatic interactions beyond 1.0 nm, and we used a 1.0 nm cutoff for van der Waals interactions. The simulation time step was set to 2 fs, and coordinates were saved every 10 ps. The temperature was maintained at 300 K using the v-rescale thermostat, while pressure was controlled using the Parrinello-Rahman barostat^111^. All simulations were performed on a single node utilizing 32 CPUs or one GPU. For each system, we discarded the first 50-ns of production MD simulations and quantitatively analyzed the subsequent 5 μs. We performed replicate simulations by extracting two random conformations from the first 50-ns of production MD, with a different seed assigned for the random initial velocities that were assigned from a Boltzmann distribution. This yielded a total of three replicates for each system, totaling 15 microseconds of production MD per system. In the first simulation, H38 was in its neutral state and a proton was placed on the ND1 position. In the second simulation H38 contained two protons at the ND1 and NE1 positions, creating the cationic form. Overall charge neutrality was maintained in both simulations through the addition of counter ions.

### Cryo-Electron Microscopy

The grids were prepared and vitrified at the Cryo-Electron Microscopy Platform (CEMP) of Helmholtz Munich. For cryo-EM grid preparation, 4.5 µL of the C9^CARD^ sample was applied to a glow-discharged 200 mesh Quantifoil R2/1 grid, blotted for 4 s with force 4 in a Vitrobot Mark III (Thermo Fisher) at 100% humidity and 4 °C. Grids were plunge frozen in liquid ethane-propane mixture, cooled by liquid nitrogen. Cryo-EM data acquisition was performed with a 300-kV, FEI Titan Krios transmission electron microscope (Thermo Fisher) with EPU software. Movie frames were recorded at a magnification of 130,000x using a Falcon4i direct electron detector camera. The total electron dose of about 55 electrons per Å^2^ was distributed over 30 frames at two different pixel sizes of 0.76 Å and 0.95 Å. The two collected datasets were processed separately and merged after extracting them to respective box sizes corresponding to a pixel size of 0.95 Å. Movies were recorded in a defocus range of −0.75 to −2.5 μm. Data processing was performed using the CryoSparc v4.4.1 helical reconstruction pipeline^112^. The final resolution between half maps and map versus model was determined after post-processing in RELION ^113^, using the Gold Standard Fourier Shell Correlation (GS-FSC) threshold value of 0.143 and 0.5, respectively. Sharpening was done with an ad-hoc B-factor of 90 Å^2^, which was selected based on the initial 3D refinement observations. Interfacial residues and surface area calculations were performed using PDBePISA (https://www.ebi.ac.uk/pdbe/pisa/). The final structural rendering was completed using UCSF Chimera (20) and UCSF ChimeraX (21).

### Pairwise sequence alignments

The C1^CARD^ (residues 1-96) and C9^CARD^ (residues 1-96) were subjected to a pairwise sequence alignment with the EMBOSS Needle webserver ^114^. This program performs a global sequence alignment using the Needleman-Wunsch algorithm^115^ and uses a default gap opening penalty of -10, a gap extension penalty of -0.5, with the alignment scored using the BLOSUM62 substitution matrix. This produced an alignment with 26 matches and 12 gaps, corresponding to a pairwise sequence identity of 27.1%.

However, the default parameters in EMBOSS Needle yielded a relatively large number of gaps (12) relative to the lengths of the C1^CARD^ and C9^CARD^ sequences (96 residues), as well as the overall structural similarity of the two domains (< 2 Å RMSD), prompted us to revisit the pairwise alignment. Based on previous work by Elofsson ^116^, we modified the gap opening (-15) and gap extension penalties (-1), which were reported to produce better global alignments with the BLOSUM62 substitution matrix. Indeed, these modifications introduced only two gaps in the alignment of C9^CARD^ and C1^CARD^ and produced a pairwise sequence identity of 19.8%. We implemented this in BioPython ^117^ version 1.85 and could produce highly similar alignments in the online EMBOSS Needle webserver when the corresponding parameters were updated (Gap Open = 15, End Gap Open = 15, End Gap = True, Gap Extend = 1, End Gap Extend = 1). Thus, we used these parameters in BioPython for the pairwise sequence alignment. For random sampling, we randomly selected *n* = 9 residues 10,000 times from the set of all surface residues (defined below), and computed the identity at these sites in the alignment. The mean and standard deviation of the percent identity values were then computed an compared to the observed value of 56%.

### Relative solvent accessible surface area

We used DSSP^118^ version 4.4.7 to calculate the relative solvent accessible surface area (rSASA) for each residue in a PDB file. The BioPython ^117^ DSSP wrapper was used, and we then parsed the results into an in-house Python script for downstream analysis. We supplied the PDB files 4rhw/chain E (C9^CARD^) and 5fna/chain A (C1^CARD^) for these calculations. Because the PDB file 5fna only contains structural data for residues 2-86 of the C1^CARD^, including an incomplete helix-6 that may produce non-realistic rSASA values, we also ran DSSP on the corresponding AlphaFold2^119^ structural model of the C1^CARD^ residues 1-96 that we obtained from the AlphaFold Database^120^ without further modification. The results presented include values derived from the AlphaFold2 model. We used a threshold of 25%, as used previously^121^, to define surface from buried residues. The sequence identity of surface and buried residues was then determined from their positions in the global pairwise sequence alignment (described above).

### ConSurf conservation scores

We extracted ConSurf conservation scores from the ConSurf-DB^81^ for PDB entries 4rhw, chain E (C9) and 5fna (C1). On the webserver, the reported grades (1-9) are derived from the raw ConSurf scores, with negative scores corresponding to more conserved positions. The assignment of grades can be extracted from the results summary (e.g. https://consurfdb.tau.ac.il/DB_NEW/4RHW/E/4RHW_E_consurf_grades.txt), and we report both values for ease of comparison to the webserver. Raw ConSurf scores for PDB 5fna are reported for residues 2-86, as these residues that are present in the PDB file.

### C9 orthologs

To examine the variation in residue type at this position of the CARD across different C9 orthologs, we extracted 182 caspase-9 sequences from the CaspBase^75^ database. We then further supplemented this dataset with an HHblits^122^ search that returned over 500 sequence hits, which were redundancy reduced with MMseqs2^123^ to identify clusters with maximum pairwise identity of 90% (319 clusters).

## Competing Interest

The authors declare no competing interests.

## Acknowledgements

We thank Prof. Lewis E. Kay (University of Toronto) for his support and insightful discussions. This study made use of NMRbox: National Center for Biomolecular NMR Data Processing and Analysis, a Biomedical Technology Research Resource (BTRR), which is supported by NIH grant P41GM111135 (NIGMS). S.R. was trained within the frame of the PhD program BioMolStruct and received funding from the Marietta Blau-Stipendium of the Austrian Agency for Education and Internationalization. We thank the Helmholtz Munich Digital Transformation and IT Department (DigIT) team for support of computational resources. T.R.A. acknowledges funding from the Initiative and Networking Fund of the Helmholtz Association (Helmholtz Investigator Grant VH-NG-20-14).

## Author Contributions

T.R.A. designed the constructs and plasmids. S.R. and T.R.A. expressed and purified protein samples. S.R. performed DSF thermal melts and saturation concentration experiments. S.R., T.M., and T.R.A collected NMR data. T.R.A processed and assigned the NMR assignment data. T.R.A processed and analyzed the NMR spin-relaxation datasets. S.R. and T.M. performed the p*K*_a_ measurement. S.R., D.K./S.B. collected negative-stain/cryo-EM data, and S.R., S.B., D.K., T. P.-K., and A.D. analyzed EM data, with S.R. and A.D. performing the processing and structure calculations. T.R.A. produced and analyzed the molecular dynamics simulations. Ð.K., I.P., and T.R.A. performed the structural bioinformatics and pairwise sequence alignments. T.R.A. conceived and designed the study, prepared the figures, and wrote the manuscript with input from S.R. All authors commented and contributed to the approved final version of the manuscript.

CRedit format:

**Swasti Rawal** – Investigation, Data Curation, Formal Analysis, Writing – Reviewing & Editing, Visualization. **Ðesika Kolarić** – Investigation, Data Curation, Formal Analysis, Writing – Reviewing & Editing. **Stefan Bohn** – Investigation, Data Curation, Writing – Reviewing & Editing. **Dagmar Kolb** – Investigation. **Tea Pavkov-Keller** – Formal Analysis, Resources, Writing – Reviewing & Editing. **Iva Pritišanac** – Investigation, Formal Analysis, Resources, Writing – Reviewing & Editing. **Tobias Madl** – Formal Analysis, Resources, Supervision, Funding Acquisition, Writing – Reviewing & Editing. **Ambroise Desfosses** – Investigation, Data Curation, Formal Analysis, Methodology, Supervision, Writing – Reviewing & Editing. **T. Reid Alderson** – Conceptualization, Investigation, Data Curation, Formal Analysis, Resources, Supervision, Project Administration, Writing – Original Draft, Writing Reviewing & Editing, Visualization, Funding Acquisition.

## Notes

### Competing Interest Statement

The authors have declared no competing interest.

