## Supplementary Information for "A histidine switch regulates pH-dependent filament formation by the caspase-9 CARD"

MDEADRRLLRRCRLRLVEEL^20^ QVDQLWDALLSRELFRPHMI^40^

EDIQRAGSGSRRDQARQLII^60^ DLETRGSQALPLFISCLEDT^80^

GQDMLASFLRTNRQAAKLS^99^

**Supplementary Figure 3**. Sequence of the C9^CARD^ residues 1-99 with the negatively (D, E) and positively charged (R, K) residues shown in red and blue, respectively. The H38 residue has been highlighted. The overall net charge at pH 7 is expected to be zero due to the equal numbers of charged residue types – 16 negative (9 D, 7 E) and 16 positive (15 R, 1 K).

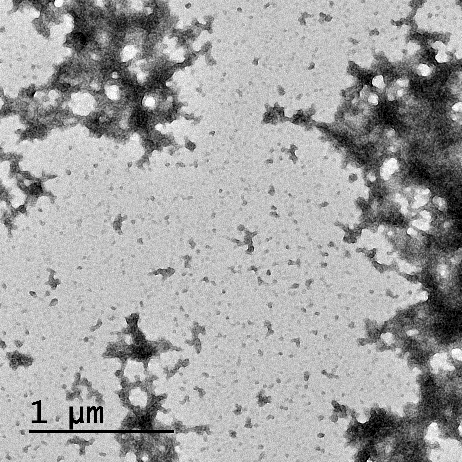

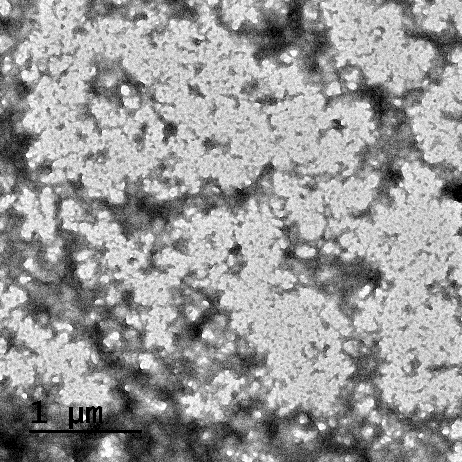

1-138	FL (C287A)

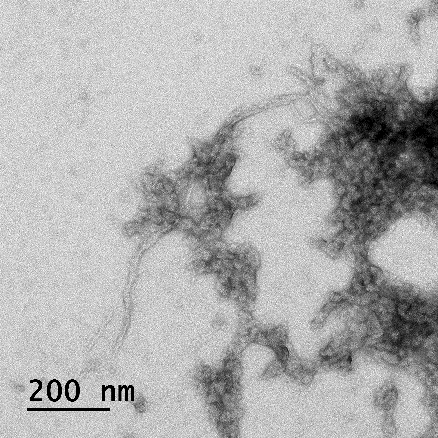

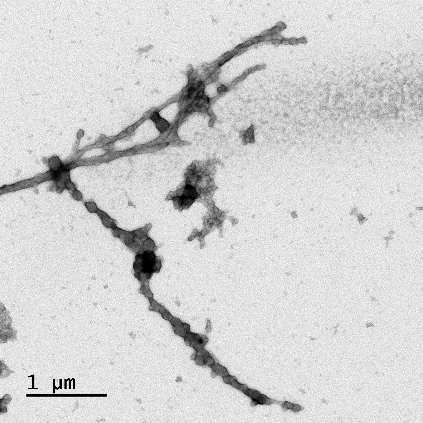

1-138	FL (C287A)

**Supplementary Figure 5**. Negative-stain EM images of C9 CARD+linker (1-138) and full-length C9 with the C287A mutation. The upper row shows representative micrographs in which amorphous aggregates are observed, which are abundant. In a few areas, filaments as observed in the bottom row, are observed. However, filaments represent a small minority of the grid.

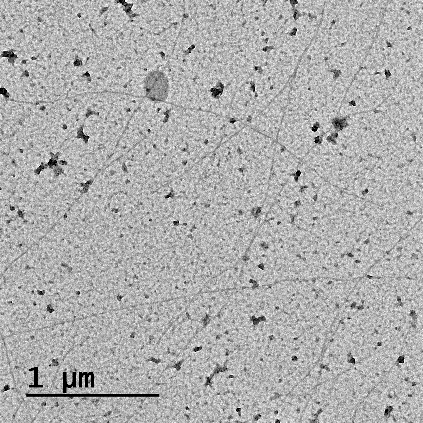

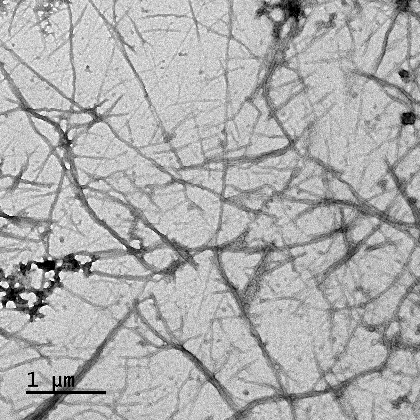

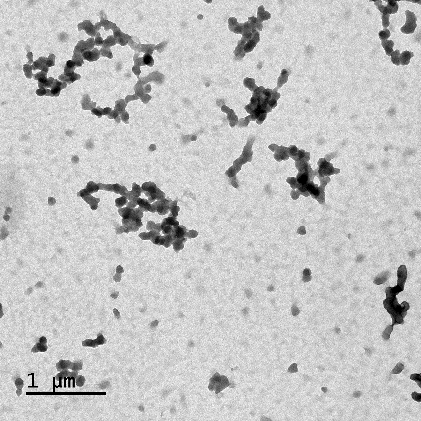

WT	H38R	H38D

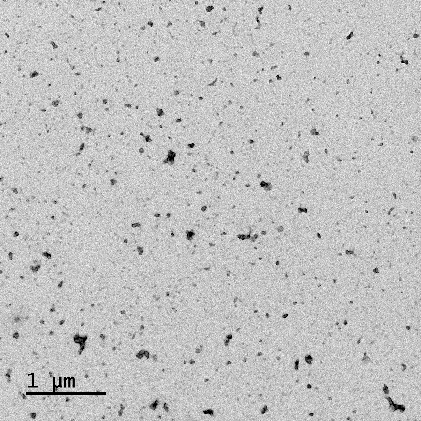

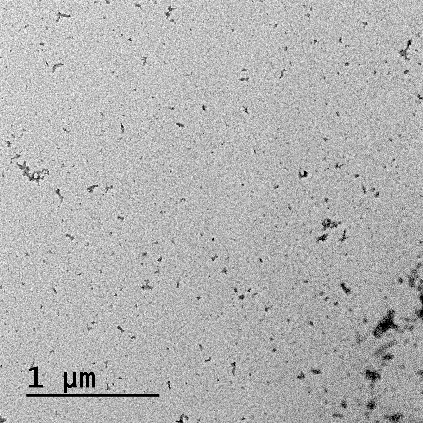

H38N	H38R, R7/R11/R56E

**Supplementary Figure 7**. Representative negative-stain micrographs of C9^CARD^ variants in pH 5.5 buffer obtained after 40 minutes of centrifugation.

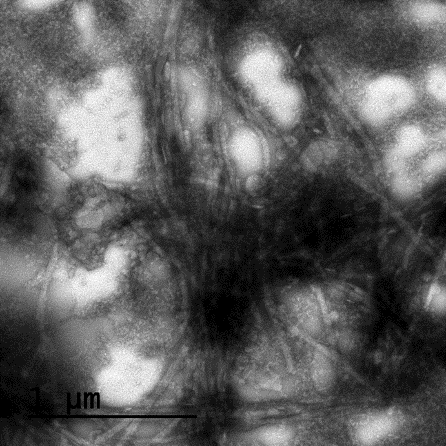

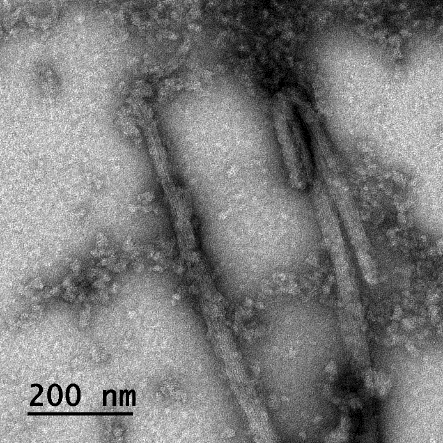

**Supplementary Figure 8**. Representative negative-stain EM micrographs of the H38D C9^CARD^ variant at pH 5.5 obtained after 80 minutes of centrifugation where some filaments are observed.

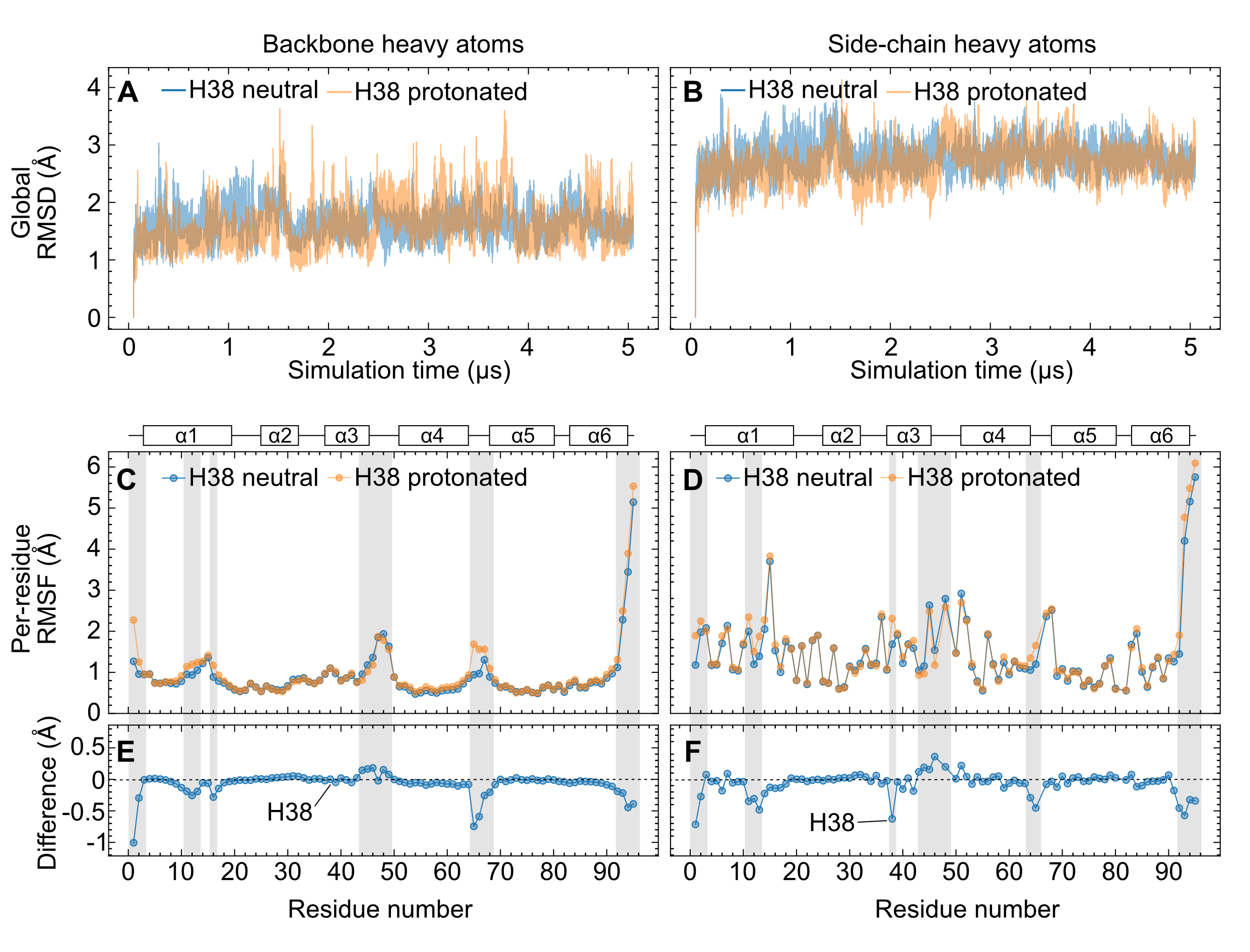

**Supplementary Figure 9. All-atom MD simulations of C9^CARD^ with either uncharged or positively charged H38.** Simulations were performed using PDB 4rwh (chain E) that was energy-minimized and equilibrated at 300 K and 1 bar. The total duration of simulation time was 5.05 μs. The initial 50 ns were discarded, and the subsequent 5 μs were analyzed here. Neutral and protonated H38 correspond, respectively, to the singly protonated N^ε2^–H τ tautomer and biprotonated N^ε2^–H and N^δ1^–H state (cationic imidazolium). Root-mean-square deviation (RMSD) values of backbone (**A**) and side-chain (**B**) heavy atoms are shown as a function of the simulation time. In both protonation states of H38, the overall fold of the protein is stable with only local structural fluctuations. Per-residue root-mean-square fluctuation (RMSF) values are shown in (**C**) and (**D**) for the backbone and side-chain heavy atoms, respectively. The differences, H38 neutral – H38 protonated, for the backbone and side-chain heavy atoms are shown in (**E**) and (**F**), respectively, with negative values corresponding to higher RMSF values in the protonated state. Residues with significant changes to their RMSF values upon H38 protonation are indicated with grey boxes. This includes the side chain of H38 itself (**F**). The most significant changes to the backbone localize to the N-terminus (M1, D2), the C-terminal region of α3 and the α3-α4 loop (Q44, R45, A46, S48), and the α4-α5 loop (R65, G66, S67, Q68). The network of interactions involving H38-M39-R65/G66-M1 is sensitive to H38 protonation. Regions of secondary structure are indicated above panels **C** and **D** for clarity.
